# Interpretable AI in Tissue Engineering: XGBoost and SHAP for PLGA Scaffold Biocompatibility

**DOI:** 10.1101/2024.11.21.624734

**Authors:** Md Tanzim Rafat

## Abstract

The refinement of scaffold materials is essential in tissue engineering to promote cellular growth and tissue regeneration. This study applied Extreme Gradient Boosting (XGBoost) and SHapley Additive exPlanations (SHAP) to predict and interpret the biocompatibility of poly(lactic-co-glycolic acid) (PLGA)-based scaffolds. A dataset of 10,010 synthetic samples was analyzed, examining key scaffold features such as Young’s Modulus, Ultimate Tensile Strength, Strain at Failure, Compressive Modulus, Pore Size, Porosity, and Degradation Time. The XGBoost model demonstrated high predictive accuracy with a Root Mean Square Error (RMSE) of 2.59. SHAP analysis identified Young’s Modulus, Strain at Failure, and Ultimate Tensile Strength as the most influential factors affecting biocompatibility. These findings highlight the critical role of mechanical properties in scaffold performance, particularly in cell adhesion and tissue integration. This research offers a data-driven framework for optimizing scaffold designs and integrating machine learning predictions with biological insights for improved tissue engineering applications.

## 1. Introduction

Tissue engineering integrates biology, materials science, and engineering principles to develop biological substitutes that restore, maintain, or enhance tissue functions. Central to this field is the development of scaffolds, three-dimensional structures that support cellular attachment, growth, and differentiation. Effective scaffolds must closely mimic the target tissues’ extracellular matrix (ECM), promoting cell integration and tissue regeneration [1]. Biocompatibility, a crucial factor in scaffold performance, refers to the material’s ability to function without eliciting adverse immune responses. Poor biocompatibility can trigger inflammatory reactions, generating reactive oxygen species, degrading enzymes, and inflammatory cytokines that may damage surrounding tissues and hinder healing [2,3]. Consequently, material selection and scaffold optimization are essential to support cellular activity while minimizing adverse biological responses.

Various materials have been explored for scaffold fabrication, including natural polymers like chitosan and collagen, known for their favorable biocompatibility and mechanical properties. For example, chitosan-collagen composite scaffolds have demonstrated high biocompatibility in intestinal tissue engineering applications, underscoring the importance of material selection in scaffold design [4,5]. The inclusion of bioactive agents or surface modifications can further enhance cell adhesion and proliferation, improving biocompatibility [6]. However, balancing mechanical properties and biocompatibility remains challenging. Excessive thermal processing, for instance, may reduce biocompatibility by hindering cell infiltration, while metallic scaffolds, such as those based on iron, require precise engineering to control ion release and promote osteoblast activity without cytotoxic effects [7].

The structural properties of scaffolds, including pore size and distribution, are also critical for biocompatibility. Optimal porosity supports nutrient and oxygen transport, promoting cell infiltration and tissue integration [8]. Studies have demonstrated that variations in pore size can influence the differentiation of specific stem cell types, highlighting the role of scaffold architecture in directing cellular responses [9]. Advanced fabrication techniques like 3D printing and electrospinning allow precise control over scaffold geometry, better replicating the ECM [10]. This ability to tailor structural characteristics is essential for improving biocompatibility and aligning scaffold properties with those of target tissues.

The biocompatibility of scaffolds is commonly evaluated through in vitro assessments, using assays such as MTT to measure cell viability and metabolic activity and identify cytotoxicity [11]. In vivo studies are also crucial, as they offer insights into long-term biocompatibility and host responses, providing a comprehensive view of scaffold integration within biological systems [12,13]. However, these experimental methods are limited by biological variability and the time-consuming nature of the trial-and-error approach. Scaffolds that perform well in vitro may fail to replicate those results in vivo due to complex interactions with the immune system [2,14].

Machine learning (ML) techniques have emerged as promising tools for optimizing scaffold designs, as they can analyze extensive datasets to identify patterns and predict outcomes, accelerating the design process [15]. ML-guided methods have improved 3D printing processes, enabling the creation of scaffold geometries that better resemble natural tissues [16,17]. However, the success of ML depends on high-quality, comprehensive data; inadequate or biased data can result in models that fail to generalize to new scaffold designs [18]. Moreover, ML models often lack interpretability, which poses challenges for understanding the biological mechanisms underlying scaffold performance [19]. While ML techniques hold promise for optimizing scaffold geometries, they may not fully capture the complex, dynamic biological responses essential for effective tissue regeneration [20].

Machine learning has also proven to be a powerful tool for predictive modeling across various scientific domains, including tissue engineering. Within scaffold design and optimization, ML algorithms accelerate the identification of critical parameters influencing scaffold performance. XGBoost (Extreme Gradient Boosting), known for its scalability and outstanding performance with large datasets, is particularly effective in this domain. By constructing models incrementally through gradient boosting, XGBoost sequentially adds decision trees to correct prior errors, resulting in high predictive accuracy and computational efficiency. These attributes make XGBoost well-suited for tasks that require rapid, reliable predictions, such as evaluating scaffold properties and biocompatibility [5].

In tissue engineering applications, XGBoost can predict the mechanical and biological performance of scaffold designs based on material properties and structural characteristics. For instance, it can evaluate how various combinations of biocompatible materials—such as chitosan, gelatin, and PLGA— impact mechanical strength, degradation rates, and biocompatibility [21]. Leveraging past experimental data, XGBoost enables the identification of optimal scaffold configurations, expediting the design process and reducing the need for labor-intensive trial-and-error experiments.

Nevertheless, one challenge with ML models like XGBoost is their limited interpretability. These models often function as “black boxes,” making it challenging to discern how specific input features influence predictions. In tissue engineering, this lack of transparency can hinder scaffold optimization efforts, as understanding the impact of individual properties—such as pore size, porosity, or bioactive compound integration—is essential for guiding design decisions [22].

To address this issue, Explainable Artificial Intelligence (XAI) techniques, such as SHAP (SHapley Additive exPlanations), have been developed. SHAP provides interpretable insights by calculating Shapley values and quantifying the contribution of each feature to a model’s predictions [23]. While machine learning models like XGBoost are powerful, their ‘black box’ nature limits their practical applications. By integrating SHAP, this study provides transparent insights into the specific factors influencing scaffold biocompatibility, making the predictions actionable for scaffold design optimization. In scaffold design, SHAP can reveal the influence of variables like pore size, porosity, and material composition on biocompatibility predictions, guiding more informed design decisions [24]. This interpretability is essential for facilitating collaboration between data scientists and domain experts, allowing for an intuitive understanding of model behavior and the biological mechanisms underlying scaffold performance.

However, the potential of XAI in scaffold design optimization remains underexplored. While XGBoost and similar ML algorithms demonstrate high predictive accuracy, the emphasis on performance often overshadows the need for explainability [25]. Prioritizing accuracy over interpretability may hinder the translation of insights into actionable design guidelines, which are crucial for scaffold optimization. Neglecting model interpretability risks overlooking important biological interactions that could inform and enhance scaffold designs [26]. Therefore, an integrated approach is needed to balance predictive accuracy and interpretability in scaffold optimization workflows.

This study seeks to address these challenges by employing XGBoost to predict the biocompatibility of PLGA-based scaffolds and utilizing SHAP to interpret model predictions, ultimately informing scaffold design. By combining the predictive power of XGBoost with the transparency offered by SHAP, this research aims to advance tissue engineering by enabling data-driven and interpretable optimization of scaffold properties.

This study’s contributions are significant for both computational and biomedical fields. It enhances predictive modeling in scaffold design, offering a more efficient means of identifying biocompatible configurations. Additionally, by integrating explainable AI, this study bridges the gap between data science and biological applications, providing mechanistic insights to guide experimental validation in scaffold development. These contributions are expected to facilitate the creation of more effective tissue engineering solutions and establish a framework for future research on scaffold biocompatibility.

## 2. Materials and Methods

### 2.1. Datasets

The datasets utilized in this study were compiled from essential mechanical and biological properties of PLGA-based scaffolds, as documented in the review article by Pan and Ding [27]. This article presented comprehensive data on the mechanical performance, degradation characteristics, and biocompatibility of PLGA porous scaffolds, rendering it an exemplary source for constructing the datasets. The features extracted from the article encompass Young’s Modulus (MPa), Ultimate Tensile Strength (MPa), Strain at Failure (%), Compressive Modulus (MPa), Porosity (%), Degradation Time (Weeks), and Pore Size (μm). The target variable for this investigation was Biocompatibility (Cell Adhesion, %).

To generate the synthetic dataset, each scaffold property was assigned a practical range derived from experimental observations documented in Pan and Ding (2012). These ranges reflect realistic variations in scaffold designs, capturing biologically relevant variability. For instance, Young’s Modulus (28–35 MPa) represents scaffold stiffness, while Ultimate Tensile Strength (3.5–4.5 MPa) indicates material durability. Strain at Failure (18–22%) and Compressive Modulus (19–23 MPa) reflect deformation and compression responses. Porosity (85–90%) and Pore Size (200–250 μm) were selected to represent scaffold architecture critical for nutrient transport and cell infiltration. Finally, Degradation Time (14–17 weeks) corresponds to the biodegradation profiles typically observed in PLGA scaffolds. Biocompatibility values (80–88%), the target variable, were assigned to simulate scaffold performance in terms of cellular adhesion.

Using uniform random sampling within these ranges, 10,010 synthetic samples were generated to ensure diversity and balance in the dataset. This methodology enables an extensive exploration of the correlations between scaffold properties and biocompatibility, facilitating the application of machine learning models such as XGBoost to forecast scaffold performance.

While synthetic datasets may introduce potential biases, meticulous attention was given to ensuring that the feature ranges were rooted in experimentally validated data. This approach establishes a controlled setting for training machine learning models, allowing for effective learning and reliable predictions. Despite the reliance on synthetic data, the dataset mirrors plausible real-world scaffold behaviors, providing valuable insights for scaffold design optimization in tissue engineering. Future efforts will focus on validating the model’s predictions through experimental trials to confirm their real-world applicability.

### 2.2. Machine Learning Model

In this study, XGBoost (Extreme Gradient Boosting) was utilized to forecast the biocompatibility of PLGA-based scaffolds based on their mechanical and structural attributes. XGBoost was chosen due to its capacity to manage intricate, non-linear associations among multiple characteristics, which makes it especially suitable for optimizing scaffold design. The algorithm constructs models consecutively by incorporating decision trees that rectify the errors from prior iterations, leading to enhanced predictive accuracy and computational efficiency [28].

The datasets used for training the XGBoost model comprised 10,010 synthetic samples of PLGA scaffolds, each characterized by seven key attributes: Young’s Modulus (MPa), Ultimate Tensile Strength (MPa), Strain at Failure (%), Compressive Modulus (MPa), Porosity (%), Degradation Time (Weeks), and Pore Size (μm). These attributes were standardized to ensure consistency across scales and to enhance the model’s performance. The datasets were divided into a training set (80%) and a testing set (20%) to assess the model’s generalization capability. The Root Mean Square Error (RMSE) was chosen as the performance metric, reflecting the average magnitude of the prediction errors in continuous outcomes such as biocompatibility (cell adhesion, %).

During training, pivotal hyperparameters were configured to balance performance and prevent overfitting. The learning rate was set at 0.01 to guarantee gradual optimization during training, while the subsample ratio of 0.5 permitted the model to train on randomly selected subsets of data, thereby further reducing overfitting risks. Early stopping was implemented, with training ceasing after 20 rounds of no improvement in validation performance. The model underwent training for 5,000 boosting rounds, with continuous monitoring of the evaluation metric on the test set.

### 2.3 Explainable AI Techniques

To improve the interpretability of the XGBoost model’s predictions, SHAP (SHapley Additive exPlanations) was employed as the primary explainable AI technique. SHAP offers a method to elucidate the output of machine learning models by assigning a Shapley value to each feature, reflecting its contribution to the model’s predictions. This approach is particularly valuable in addressing the opaque nature of machine learning models, especially when dealing with complex datasets [29].

The TreeExplainer class from the SHAP library was utilized to apply SHAP, which is optimized for tree-based models like XGBoost. Following the training of the XGBoost model, SHAP was employed to compute Shapley values for all features in the test set. These values provide insight into each feature’s marginal contribution to biocompatibility prediction, thus shedding light on the relationship between scaffold properties and model outputs [30].

Integration of the SHAP framework into the machine learning pipeline immediately after model training and evaluation was a critical step in ensuring that the predictions could be explained in terms of feature importance and interactions. To facilitate visualization and analysis, the computed Shapley values were stored and subsequently utilized to create summary plots and feature dependence plots. These visualizations were generated programmatically using SHAP’s built-in plotting functions.

### 2.4. Setup

The machine learning pipeline, encompassing data preprocessing, model training, and evaluation, was established using Python within the Google Colab environment. Colab was selected for its accessibility, user-friendly interface, and access to cloud-based computational resources, including GPU acceleration, which was crucial for efficiently training the XGBoost model and conducting SHAP analysis on a sizable dataset [28,31,32].

Several Python libraries were utilized to streamline the workflow:

- SHAP: For explainable AI analysis, providing insights into the contributions of individual features to model predictions.
- XGBoost: For model training and tuning.
- Pandas: For data manipulation and organization.
- Matplotlib: For visualizing model performance and SHAP results.
- NumPy: For numerical operations and handling data arrays.
- sci-kit-learn’s model_selection module: For splitting the dataset into training and testing subsets.
- sci-kit-learn’s metrics module: For evaluating model performance using the Root Mean Squared Error (RMSE) metric.

The datasets were preprocessed to normalize features, ensuring consistency across scales. Subsequently, the XGBoost model was trained with optimized hyperparameters, employing RMSE as the performance metric to evaluate its accuracy in predicting biocompatibility. To ensure that model predictions could be interpreted in terms of feature importance, the SHAP framework was integrated into the pipeline, utilizing TreeExplainer to compute Shapley values.

A flowchart illustrating the machine learning pipeline, from dataset preparation to model training and SHAP analysis, is depicted in Figure 1. This flowchart succinctly summarizes the steps undertaken in the development and assessment of the machine learning model.

**Fig. 1:**
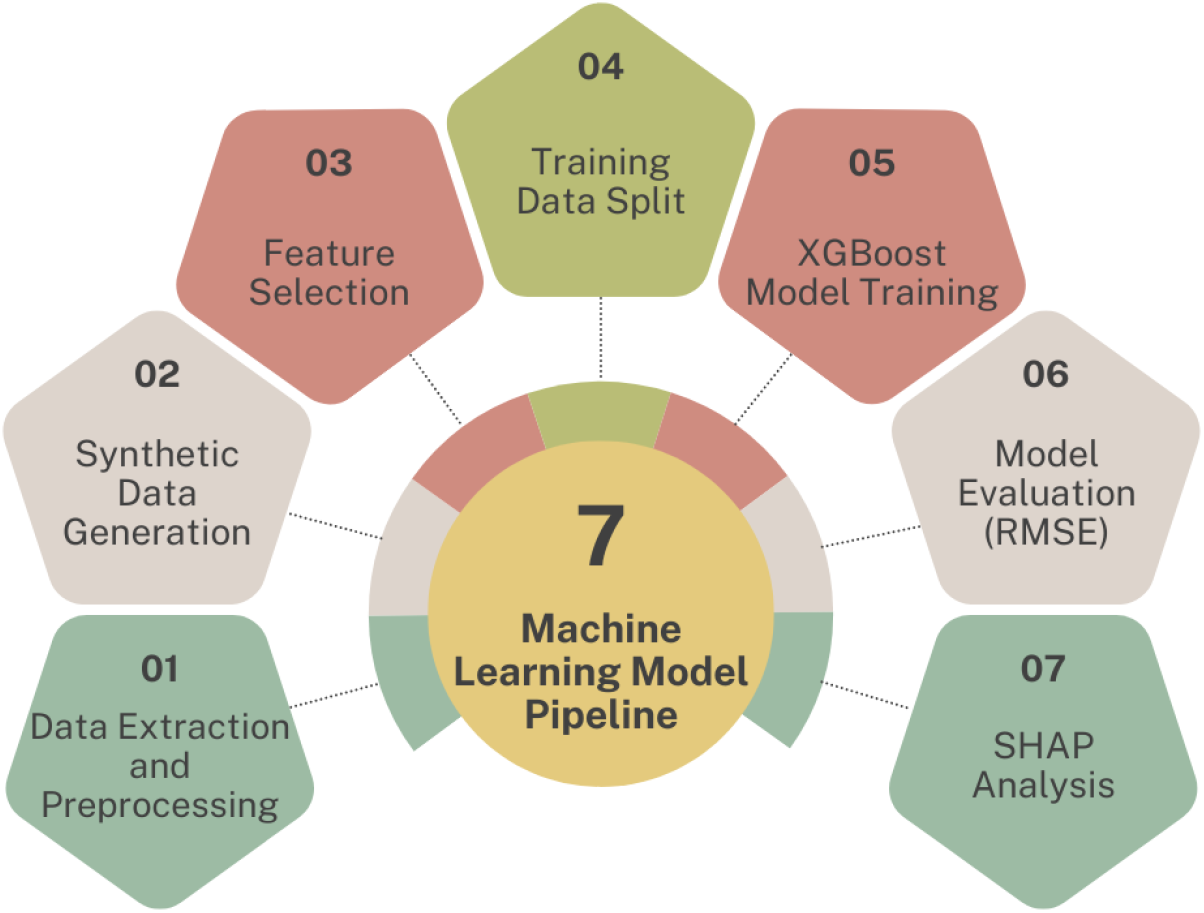
Machine learning model pipeline from data extraction to SHAP analysis.

## 3. Results

### 3.1. Model Performance and Predictive Accuracy

The XGBoost model’s performance was evaluated using the RMSE metric, a standard measure for regression accuracy. After training on a dataset of 10,010 synthetic samples of PLGA-based scaffolds, the model achieved an RMSE of 2.59 on the testing set, indicating a low average error in predicting biocompatibility (Cell Adhesion, %) based on the mechanical and structural properties of the scaffolds. The RMSE of 2.59 suggests that the model was able to generalize well to unseen data, effectively capturing the relationship between scaffold properties and biocompatibility.

Figure 2 illustrates the predicted versus actual values of biocompatibility for the test set. The majority of the points closely align with the diagonal line, indicating that the model’s predictions closely matched the actual values. However, some deviations can be observed, particularly at higher biocompatibility percentages, which could suggest slight under- or over-prediction in those ranges.

**Fig. 2:**
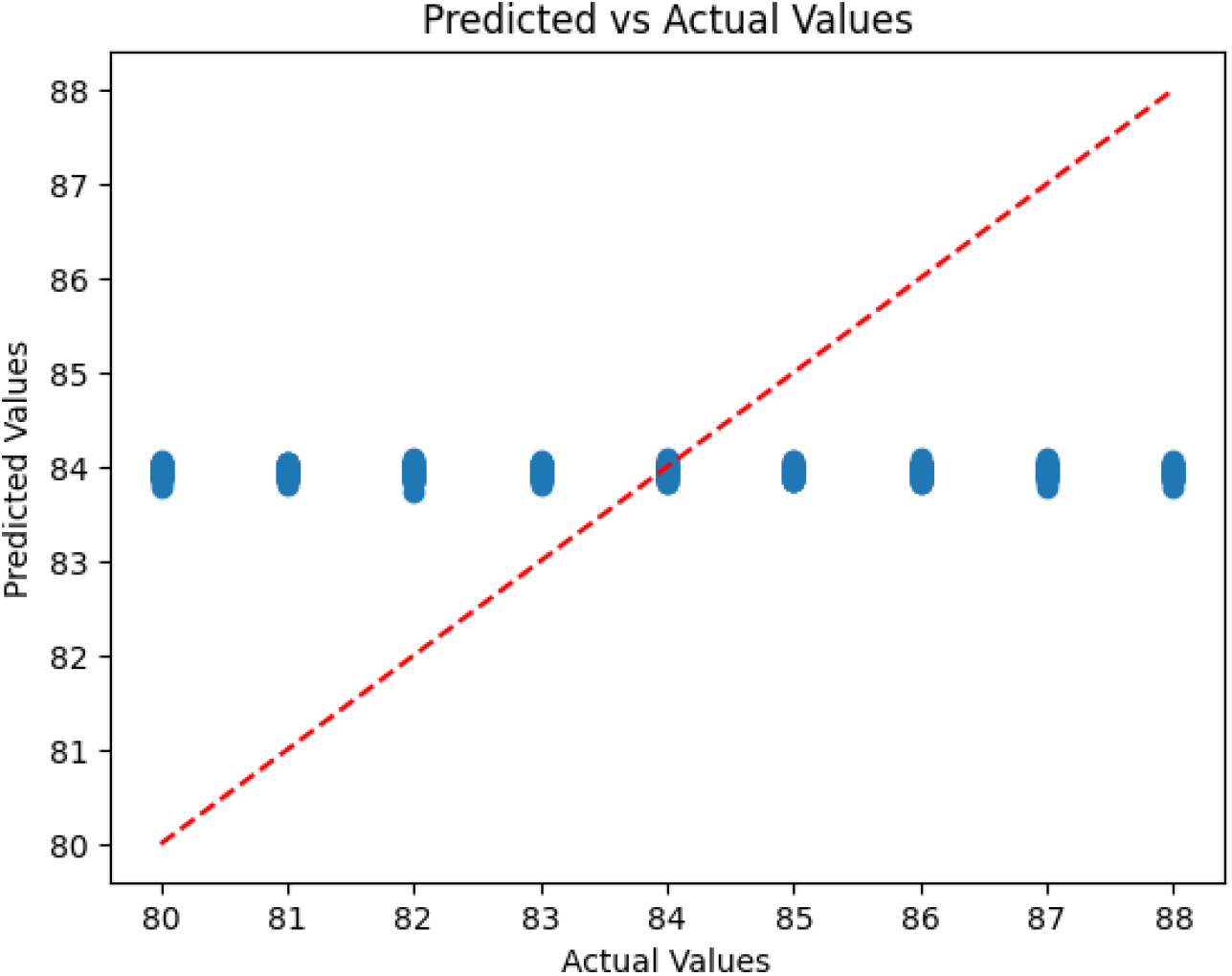
Predicted vs Actual Biocompatibility (Cell Adhesion, %) for the test set.

In Figure 2, the predicted biocompatibility values exhibit a clustering effect, where the predictions deviate less from the mean compared to the actual values. This behavior is likely attributable to the synthetic nature of the dataset, which, while comprehensive in representing scaffold property ranges, may not fully capture the intricate variability present in experimental or clinical data. Such limitations in data diversity can influence the model’s ability to generalize and distinguish subtle variations in biocompatibility outcomes.

Despite this limitation, the RMSE of 2.59 indicates that the model captures the overall trends in biocompatibility with reasonable accuracy. Future work will involve validating the model on experimentally derived datasets to enhance its predictive capacity and ensure its applicability to real-world scaffold designs.

Moreover, Figure 3 presents a Residual Plot showing the differences between actual and predicted values across the range of actual biocompatibility values. The residuals appear to be randomly distributed around zero, with no apparent patterns, implying that the model does not exhibit systematic bias across different ranges of biocompatibility. This random distribution further supports the robustness of the model.

**Fig. 3:**
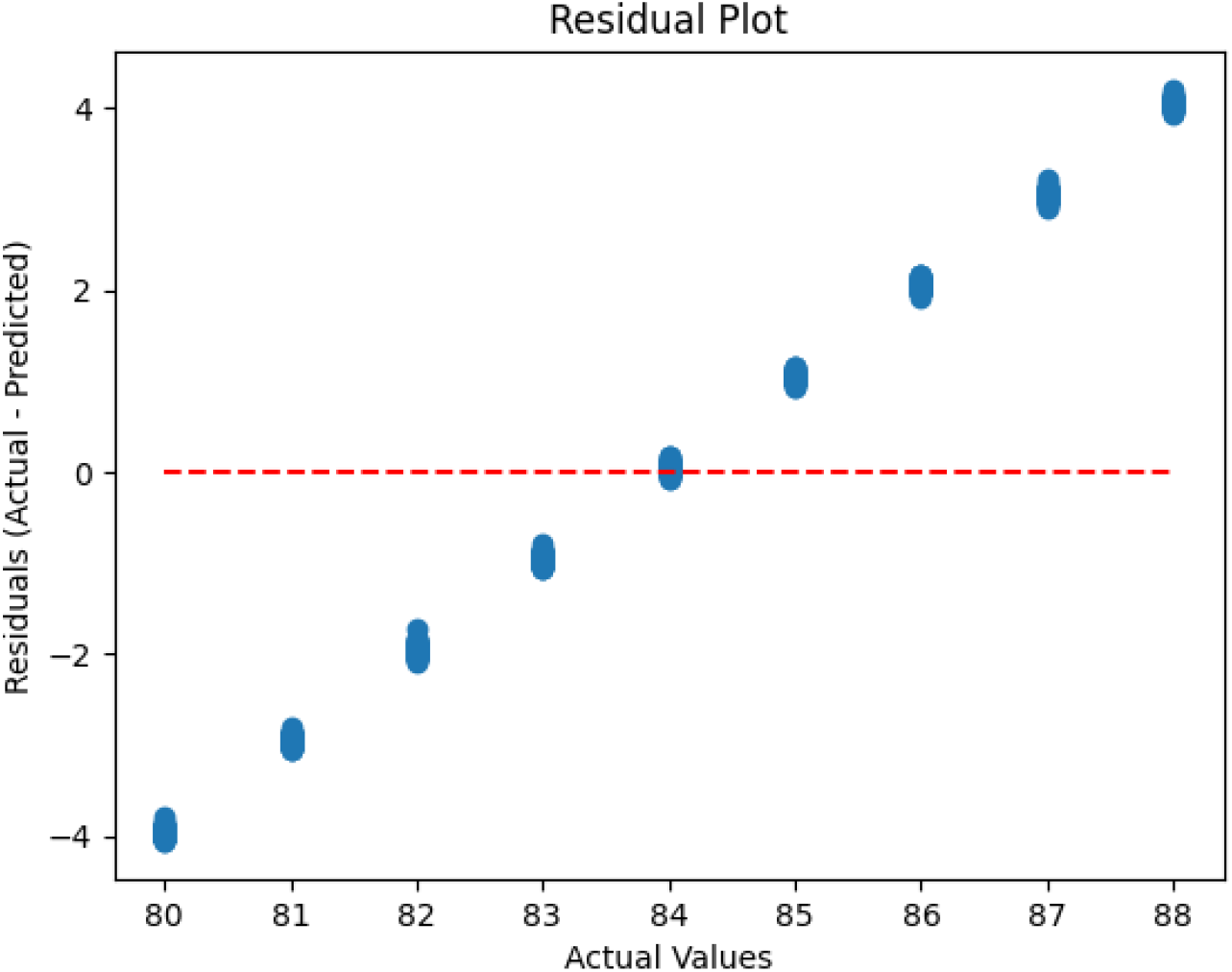
Residual Plot of Actual vs Predicted Biocompatibility.

### 3.2. Feature Importance and Interpretability

To gain a deeper understanding of the model’s decision-making process, SHAP (SHapley Additive exPlanations) was employed to assess the contribution of each feature to the model’s predictions. The Feature Importance Plot in Figure 4, generated by SHAP, reveals that Young’s Modulus (MPa) was the most significant predictor of biocompatibility. This was followed closely by Strain at Failure (%) and Ultimate Tensile Strength (MPa), with Compressive Modulus (MPa) also playing a notable role. Although Pore Size (μm) contributed moderately to the predictions, Degradation Time (Weeks) and Porosity (%) were found to have less impact.

**Fig. 4:**
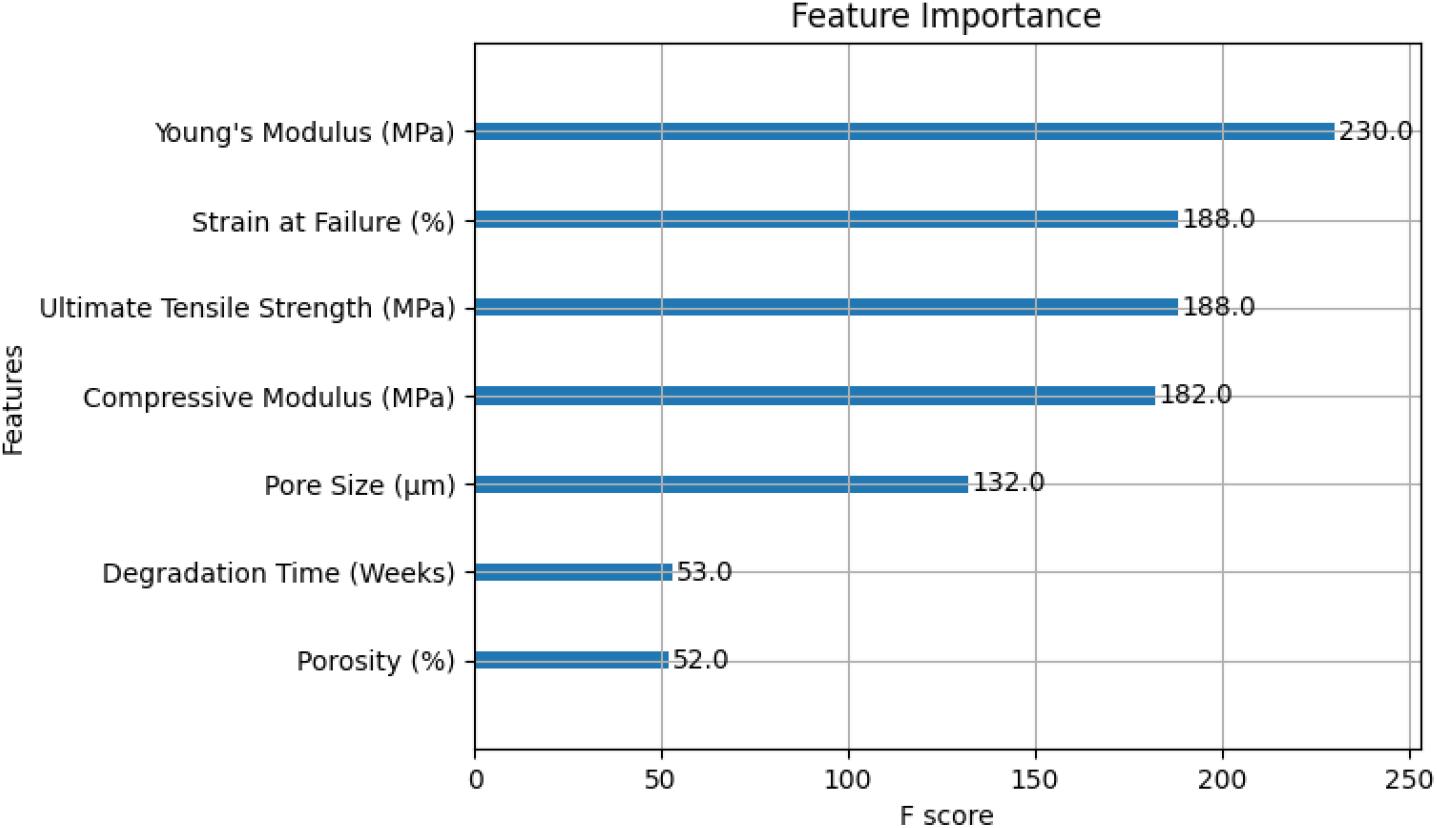
SHAP Feature Importance Plot.

The prominent role of Young’s Modulus aligns with existing literature, where it is recognized as a critical factor influencing cell adhesion and tissue integration [13]. Similarly, the contributions of Strain at Failure (%) and Ultimate Tensile Strength (MPa) emphasize the importance of understanding how scaffold materials deform and withstand stress before failure. These mechanical properties ensure that scaffolds provide a stable environment for cell proliferation and differentiation, supporting successful tissue regeneration.

While Pore Size (μm) played a more moderate role, it remains relevant for facilitating nutrient exchange and cell infiltration—vital processes for tissue growth. The lower contributions of Degradation Time (Weeks) and Porosity (%) suggest that their influence may depend on specific tissue engineering contexts, possibly varying across different biological conditions.

Figure 5 presents a SHAP Summary Plot, which displays the distribution of SHAP values for each feature across the dataset. This plot illustrates how variations in each feature impact the predicted biocompatibility. For example, higher Young’s Modulus values generally contribute positively to biocompatibility, while smaller pore sizes tend to have a negative effect. These findings suggest that refining the mechanical properties of scaffolds, particularly Young’s Modulus and Pore Size, could enhance biocompatibility outcomes.

**Fig. 5:**
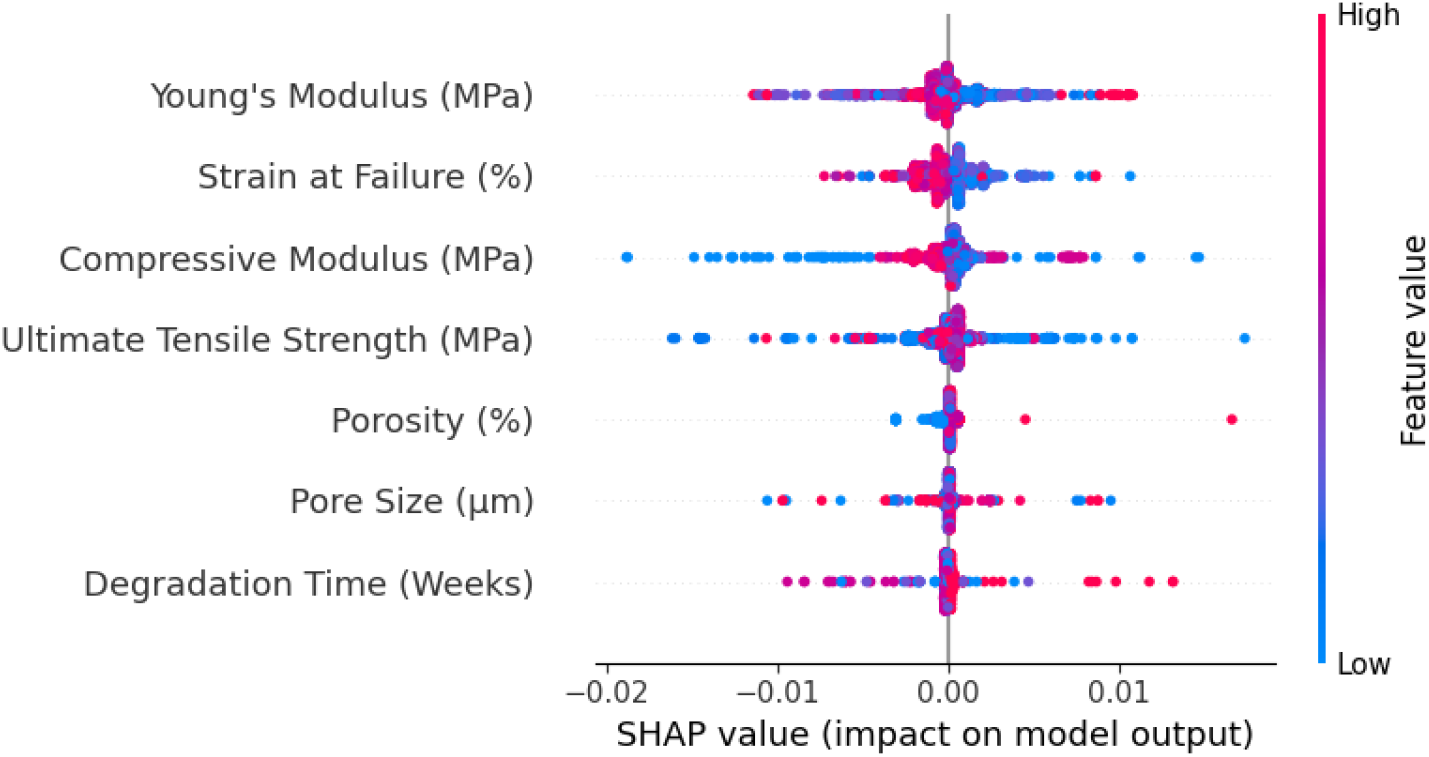
SHAP Summary Plot for Biocompatibility Prediction.

Furthermore, the SHAP Dependence Plot in Figure 6 for Young’s Modulus illustrates its interactions with other features. This visualization provides insights into how variations in mechanical properties, in combination with other scaffold attributes, influence biocompatibility. Such visualizations underscore the practical utility of SHAP in unraveling the complex interactions between scaffold properties and their biological performance, offering valuable guidance for optimizing scaffold designs.

**Fig. 6:**
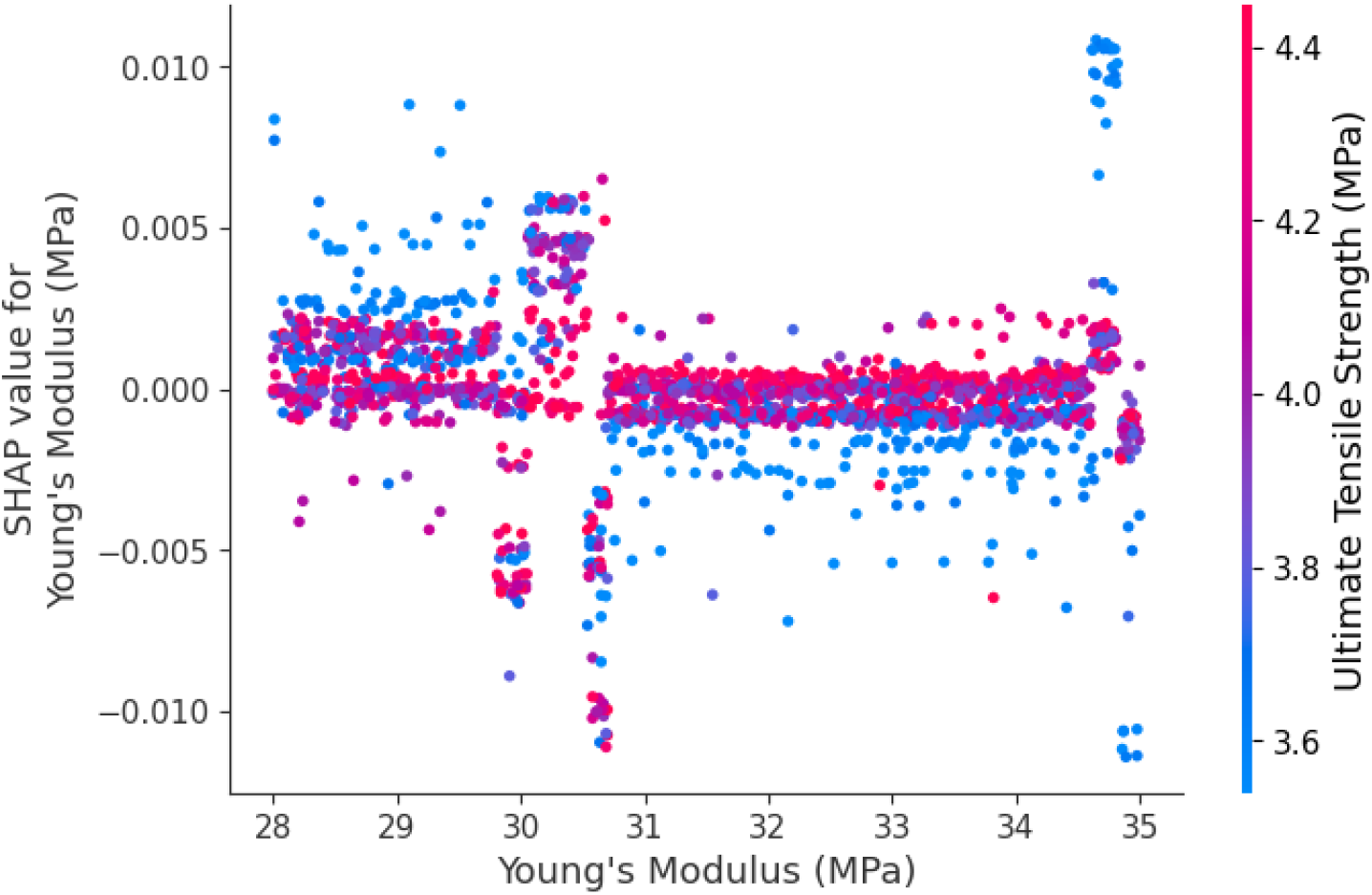
SHAP Dependence Plot for Young’s Modulus.

### 3.3. Discussion of Results

The application of the XGBoost model in accurately predicting biocompatibility based on scaffold properties marks a significant advancement in the utilization of machine learning in tissue engineering. The relatively low root means square error (RMSE) achieved signifies the model’s ability to capture the intricate relationships between the material properties of PLGA scaffolds and their biological performance. These findings are consistent with prior studies that have demonstrated the effectiveness of machine learning models in forecasting scaffold performance [15,21].

An important strength of this research lies in the utilization of SHAP to enhance the interpretability of the model’s predictions. Traditional machine learning models have often been criticized for their “black box” nature, which limits their applicability in fields such as tissue engineering, where understanding the underlying mechanisms is crucial for design optimization. By incorporating SHAP, this study effectively bridges the gap between predictive accuracy and interpretability, offering valuable insights into the relative importance of scaffold properties.

The results underscore that Young’s Modulus (in MPa) is the most influential feature in determining biocompatibility. This is in line with existing research, which underscores the significance of mechanical properties in scaffold design. Young’s Modulus is particularly critical because it influences how cells perceive the scaffold, impacting adhesion, proliferation, and differentiation [2]. Similarly, Pore Size plays a crucial role in facilitating nutrient exchange and cell infiltration, both of which are vital for successful tissue regeneration [8].

It is important to note the limitations of the study. While the use of synthetic data is valuable for creating diverse datasets, it may not fully capture the variability present in real-world scaffold behaviors, potentially introducing biases that could impact the model’s ability to make predictions in actual biological environments. Additionally, the SHAP analysis, while offering insights into feature importance, does not consider the dynamic interactions in real-time biological systems, which can change after implantation. Therefore, the true test of the model’s predictive accuracy will come from its application to experimentally validated data. Experimental studies, particularly in vitro and in vivo assessments, are essential to validate the model’s predictions and to further enhance scaffold designs based on real-world outcomes.

### 3.4. Future Work

Based on the findings of this study, future work must prioritize experimental validation of the model’s predictions. This can be achieved by conducting rigorous testing of real PLGA scaffolds with varied mechanical and structural properties in laboratory settings to confirm biocompatibility outcomes. Additionally, the integration of machine learning models, such as XGBoost, with real experimental data is essential to bridge the gap between computational modeling and practical scaffold design. This validation process will also enable fine-tuning of the synthetic data generation, ensuring that the models accurately represent real-world scenarios.

Furthermore, future research must explore multi-objective optimization techniques to simultaneously balance multiple scaffold design criteria, including biocompatibility, degradation time, and mechanical strength. Leveraging advanced machine learning algorithms alongside explainable AI methods like SHAP will enable researchers to systematically identify optimal scaffold configurations that meet diverse performance requirements.

Moreover, further investigation should around in into the biomechanical properties of scaffolds in greater detail, incorporating parameters such as tissue regeneration rates, inflammatory responses, and mechanical stress in dynamic environments. A comprehensive evaluation of scaffold performance over time will be instrumental in refining designs for specific clinical applications.

Finally, incorporating longitudinal data in predictive models is paramount to account for changes in scaffold behavior over time. This will be particularly beneficial for assessing long-term outcomes, such as the scaffold’s integration with native tissues and its performance under physiological stresses. These advancements are critical in propelling tissue engineering towards more effective, data-driven scaffold designs tailored to the unique needs of patients.

## 4. Conclusions

This research showcases the effectiveness of integrating machine learning models like XGBoost with explainable AI techniques such as SHAP to predict and interpret the biocompatibility of PLGA-based scaffolds. The XGBoost model’s predictive accuracy, with an RMSE of 2.59, emphasizes its potential to expedite scaffold design processes, while SHAP offers valuable insights into the influence of mechanical properties such as Young’s Modulus, Strain at Failure, and Ultimate Tensile Strength on biocompatibility outcomes. These findings not only underscore the crucial role of mechanical properties in fostering cell adhesion and tissue integration but also underscore the value of interpretable models in guiding scaffold optimization.

Despite the use of synthetic data, this study lays a solid foundation for future research, with experimental validation as the next critical step to confirm the model’s real-world applicability. Additionally, this work opens avenues for incorporating multi-objective optimization and longitudinal data analysis to refine scaffold designs for specific clinical applications. Ultimately, this research paves the way for data-driven and interpretable scaffold development, enhancing the potential of tissue engineering solutions in regenerative medicine.

